# *De novo* clustering of long-read transcriptome data using a greedy, quality-value based algorithm

**DOI:** 10.1101/463463

**Authors:** Kristoffer Sahlin, Paul Medvedev

## Abstract

Long-read sequencing of transcripts with PacBio Iso-Seq and Oxford Nanopore Technologies has proven to be central to the study of complex isoform landscapes in many organisms. However, current *de novo* transcript reconstruction algorithms from long-read data are limited, leaving the potential of these technologies unfulfilled. A common bottleneck is the dearth of scalable and accurate algorithms for clustering long reads according to their gene family of origin. To address this challenge, we develop isONclust, a clustering algorithm that is greedy (in order to scale) and makes use of quality values (in order to handle variable error rates). We test isONclust on three simulated and five biological datasets, across a breadth of organisms, technologies, and read depths. Our results demonstrate that isONclust is a substantial improvement over previous approaches, both in terms of overall accuracy and/or scalability to large datasets. Our tool is available at https://github.com/ksahlin/isONclust.

## 1 Introduction

Long-read sequencing of transcripts with Pacific Biosciences (PacBio) Iso-Seq and Oxford Nanopore Technologies (ONT) has proven to be central to the study of complex isoform landscapes in, *e.g.*, humans [1–4], animals [5], plants [6], fungi [7] and viruses [8]. Long reads can reconstruct more complex regions than can short RNA-seq reads because the often complex assembly step is not required. However, they suffer from higher error rates, which present different challenges. Using a reference genome can help alleviate these challenges, but, for non-model organisms or for complex gene families, *de novo* transcript reconstruction methods are required [2, 9].

For non-targeted Iso-Seq data, the commonly used tool for *de novo* transcript reconstruction is the ToFU pipeline from PacBio [7]. However, ToFU generates a large number of redundant transcripts [10, 11, 7], and most studies have had to resort to additional short-read sequencing [11, 12] or relying on a draft reference [5]. For ONT data, there are, to the best of our knowledge, no methods yet for *de novo* transcript reconstruction. Therefore, we believe that the full potential of long-read technologies for *de novo* transcript reconstruction from non-targeted data has yet to be fully realized.

Algorithms for this problem are roughly composed of two steps [2, 7]. Since most transcripts are captured in their entirety by some read, there is no need to detect dovetail overlaps between reads (as in RNA-seq assembly). Instead, the first step is to group reads together into clusters according to their origin, and the second is to error-correct the reads using the information within each cluster. This is the approach taken by ToFU, but it clusters reads according to their isoform (rather than gene) of origin. This separates reads that share exons into different clusters – reducing the effective coverage for downstream error correction. For genes with multiple isoforms, this signficantly fragments the clustering and, we suspect, causes many of the redundant transcripts that have been reported. For ONT data, there exists a tool to cluster reads into their gene family of origin (carnac-lr [9]), but it performed sub-optimally in our experiments and scales poorly for larger datasets. Thus, better clustering methods are need to realize the full potential of long reads in this setting. There exists a plethora of algorithms for *de novo* clustering of generic nucleotide-[13–16], and protein-sequences [17, 14, 18, 19]. Several algorithms have also been proposed for clustering of specific nucleotide data such as barcode sequences [20], EST sequences [21–23], full-length cDNA [24], RAD-seq [25], genomic or metagenomic short reads [26–31], UMI-tagged reads [32], full genomes and metagenomes [33], and contigs from RNA-seq assemblies [34]. However, our clustering problem has unique distinguishing characteristics: transcripts from the same gene have large indels due to alternative splicing, and the error rate and profile differs both between [2] and within [35] reads. Furthermore, the large number of reads limits the scalability of algorithms that require an all-to-all similarity computation. *De novo* clustering of long-read transcript sequences has to our knowledge only been studied in [2, 36, 7] for Iso-Seq data and in [9] for ONT data. However, neither IsoCon [2] nor Cogent [36] scale to large datasets, and the limitations of ToFu [7] and carnac-lr [9] have already been described above. In [30], the authors demonstrated that using quality values can significantly improve clustering accuracy, especially for higher error rates, but their method was not designed for long reads.

Motivated by the shortcomings of the existing tools, we develop isONclust, an algorithm for clustering long reads according to their gene family of origin. isONclust is available at https://github.com/ksahlin/isONclust. isONclust is greedy (in order to scale) and makes use of quality values (in order to handle variable error rates). We test isONclust on three simulated and five biological datasets, across a breadth of organisms, technologies, and read depths. Our results demonstrate that isONclust is a substantial improvement over previous approaches, both in terms of overall accuracy and/or scalability to large datasets. isONclust opens the door to the development of more scalable and more accurate methods for *de novo* transcript reconstruction from long-read datasets.

## 2 Methods

### 2.1 Definitions

Let *r* be a string of nucleotides that we refer to as a read. Let *q*(*r_i_*) be the probability of base call error at position 1 *≤ i ≤ |r|*. This value can be derived from the Phred quality value *Q* at position *I* as *q*(*r_i_*) = 10^*−*(*Q/*10)^. Let *ϵ_r_* be the average base error rate, 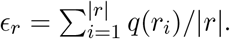 Given two integers *w* and *k* such that 1 *≤ k ≤ w ≤ |r|*, the *minimizer at position i* is the lexicographically smallest substring of *r* of length *k* that starts at a position in the interval of [*i, i* + *w*) [37]. Let *M* (*r*) be the set of ordered pairs containing all the minimizers of *r* and their start positions on the read. For example, for *r* = *ACGCCGATC, k* = 2*, w* = 4, we have *M* (*r*) = {(*AC,* 0), (*CC,* 3), (*AT,* 6)}. All the strings of *M* (*r*) are referred to as the *minimizers* of *r*.

### 2.2 **isONclust** overview

isONclust is a greedy clustering algorithm. Initially, we sort the reads so that sequences that are longer and have higher quality scores appear earlier (details in Section 2.3). We then process the reads one by one, in this sorted order. At any point, we maintain a clustering of the reads processed so far, and, for each cluster, we maintain one of the reads as the *representative* of the cluster. We also maintain a hash-table *H* such that for any *k*-mer *x*, *H*(*x*) returns all representatives that have *x* as a minimizer.

At each point that a new read is processed, isONclust consists of three steps. In the first step, we find the number of minimizers shared with each of the current representatives, by querying *H* for each minimizer in the read and maintaining an array of representative counts. We refer to any representative that shares at least one minimizer with the read as a *candidate*. In the second step, we use a minimizer-based method to try to assign a read to one of the candidate representative’s cluster. To do this, we process the candidate representatives in the order of most shared minimizers to the least. For each representative, we estimate the fraction of the read’s sequence that is shared with it (details in Section 2.3). If the fraction is above 0.7, then we assign the read to that representative’s cluster; if not, we proceed to the next candidate. However, if the number of shared minimizers with the current representative drops below 70% of the top-sharing representative, or below an absolute value of 5, we terminate the search and proceed to the third step.

In the third step, we fall back on a slower but exact Smith-Waterman alignment algorithm. If two transcripts have highly variable exon structure or contain many mutations (e.g., multi-copy genes or highly-mutated allele), then it could create long regions of no shared minimizers and prevent the minimizer matching approach from detecting the similarity. An alignment approach is more sensitive and can detect that they should be clustered together. To control the run-time, we align the read only to the representative with the most shared minimizers (several in the case of a tie). Similar to the second step, we estimate the fraction of the read’s sequence that is shared with the representative (details in Section 2.3), and if the quality is above a threshold (details in Section 2.3), the read is assigned to that representative’s cluster. Otherwise, the read is assigned to a new cluster and made its representative.

### 2.3 **isONclust** in-depth

#### Homopolymer compression

An important aspect of isONclust is that reads are homopolymer compressed, i.e. all consecutive appearances of a base are replaced by a single occurrence. For example, the sequence GCCTGGG is replaced by GCTG. When a homopolymer is compressed, the base quality assigned to the compressed base is taken as the highest quality of the bases in the original homopolymer region. The reason for using the highest quality value is that it is a lower bound on the presence of at least one nucleotide in that run. The homopolymer compression removes a large amount of homopolymer indel errors during minimizer matching or alignment and at the same time removes repetitive minimizers (from, e.g., poly-A tails). We borrowed this idea from [38, 39], where it was used to improve the sensitivity of PacBio read alignment.

#### Sorting order

Prior to greedily traversing the reads, we sort them in decreasing order of their score. We define the score *s*(*r*) of a read *r* as the expected number of error-free *k*-mers in *r*. Let *X_i_* be a binary indicator variable modelling the event that the *k*-mer starting at position *i* of *r* has no sequencing error (*X_i_* = 1). Then, we have

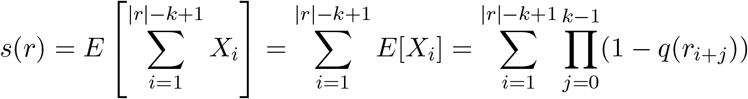

The score of a read can be quickly computed in a linear scan by maintaining a running product over a sliding window of *k* quality scores.

The ordering produced by this score function is crucial for the accuracy of our greedy approach. Observe that our algorithm never re-computes which read in a cluster is the representative, and all future reads are compared only to a cluster’s representative and not to other reads in the cluster. This is done for the sake of efficiency, but, as a downside, once a read initiates a new cluster, it becomes its representative forever. However, our score function guarantees that it will have the largest expected number of error-free *k*-mers of any future read in the cluster. In the case of alternatively spliced genes, this means that the representative will contain the most complete exon repertoire of the gene. This allows us to make the assumption that all exon differences during minimizer matching or alignment are encountered as deletions with respect to the representative. In both the matching and alignment parts, we will therefore not penalize for long deletions in the read. We do penalize for insertions in reads with respect to representatives because we assume that they cannot be due to exon differences. We note that in the cases that our assumption does not hold (e.g. when several exons are not present in the longest isoform), we may miss some matches and/or alignments. However, we do tolerate some fraction of unmatched sequence in later steps.

#### Estimating fraction of shared sequence, based on a minimizer match

Consider a read *r*, a representative *c*, the set *M* (*r*), and the minimizers of *r* that are shared with *c*. We would like to quickly estimate the fraction *f* of *r*’s sequence that would align to *c*, if an alignment were to have been performed. When two consecutive minimizers in *M* (*r*) match *c*, we simply count the sequence spanned between their positions towards *f*. For the harder case, consider a sequence of *i* consecutive unmatched minimizers in *r* that are flanked on both sides by either a matched minimizer or the end of the read. We must decide if this is due to the region being unalignable or due to true sequencing errors. Let *p*(*ϵ_r_, ϵ_c_*) be the probability, known *a priori*, that a minimizer in a read is not matched to another read, given that they are both generated from the same transcript with respective error rates *ϵ_r_* and *ϵ_c_*. Then, the probability that *i* consecutive minimizers of *r* are unmatched as a result of sequencing error can be estimated as *p*(*ϵ_r_, ϵ_c_*)^*i*^. If this probability is above 0.1, we count the whole region towards *f*, otherwise we do not.

#### Estimating *a priori* minimizer mismatch rate

Deriving an analytical formula for *p*(*ϵ_r_, ϵ_c_*) is a challenge, as it is a complex function depending on, e.g., the sequence of the true minimizer in the window, the sequence in the window, the error profile, and the properties of homopolymer compression. Instead, we use simulations to create a lookup table for *p*. We randomly generate a transcript of length 1 kbp and, from that transcript, two reads *r* and *c* with error rates *ϵ_r_* and *ϵ_c_*, respectively. The errors are equally distributed between insertion and deletion errors since Iso-Seq and ONT errors are dominated by indels. Further customization of the error profile to more accurately reflect the technology is possible, but we found that it had little effect. We then homopolymer compress the reads and count the fraction of *r*’s minimizers that do not match *c*. We repeat the process 1,000 times, each time starting with a new transcript. The average fraction of non-matching minimizers is used as the estimate for *p*(*ϵ_r_, ϵ_c_*). We pre-computed the lookup table for a range of *ϵ_r_* and *ϵ_c_* values that we observe in practice, but it can also be computed on the fly for datasets outside of these ranges.

#### Estimating fraction of shared sequence, based on the alignment

When the minimizer matching approach fails to find a match, isONclust aligns the read *r* to the most promising representative *c* using the parasail [40] implementation of Smith-Waterman. Let *ϵ* = *ϵ_r_* + *ϵ_c_* be the combined error rate of *r* and *c*. The parameters to Smith-Waterman are given in Appendix A.1 and are a function of *ϵ*. Based on this alignment, we would like to estimate the fraction of *r* whose alignment to *c* is consistent with having the same underlying sequence but allowing sequencing errors (i.e. the same goal as we had during minimizer matching). We aim to tolerate a mismatch rate of *ϵ*. Consider the pairwise alignment *A*, represented as a matrix where the two rows correspond to *r* and *c*, and each cell contains a symbol indicating either a match, mismatch, or a gap. Consider a window of *k* columns in *A* starting at position *i* of *r*. Let *W_i_* = 1 if the number of columns in the window that are not matches is ≤ *⎡Ek⎤*; otherwise, let *W_i_* = 0. We let the shared fraction 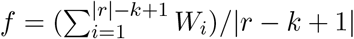 and add *r* to the cluster of *c* if *f* is above 0.4.

#### Time complexity

Our tool is a greedy heuristic, hence it is challenging to derive a worst-case run-time that is informative. We attempt to do so by parametrizing our analysis and fixing the number of representatives identified as candidates for a read as *d*. The initial sorting step takes *𝒪*(*n* log *n*) time. Then for each read, the identification of minimizers takes *𝒪*(*ℓ*) time, where *ℓ* is the read length. Here, we treat *w* and *k* as constants. There are at most *ℓ* minimizers, and each one hits at most *d* representatives, hence identifying candidate representatives takes *𝒪*(*ℓd*) time. Ranking the candidate representatives can be done using counting sort in *𝒪*(*d*) time. For minimizer matching, each of the at most *d* candidates can be processed using a linear scan through the read, leading to a total of *𝒪*(*ℓd*) time. The alignment step is done only once and is dominated by the *𝒪*(*ℓ*^2^) Smith-Waterman time. Hence, the total run-time is *𝒪*(*n* log *n* + *nℓd* + *nℓ*^2^). In the worst-case, *d* can be *Ω(n*), but it is much less in practice.

#### Parameters and thresholds

The only parameters to isONclust are the window size *w* and the *k*-mer size *k*. We found through trial-and-error that *k* = 15 and *w* = 50 work well for Iso-Seq data, and *k* = 13 and *w* = 20 work well for ONT data. Note that these lengths are applied for homopolymer compressed reads, thus a 13-mer is likely to be much longer in the original read. There are also several other hard threshold used by isONclust, as described above. We set these through a mix of intuition and testing on simulated data; nevertheless, we found that isONclust is robust to these thresholds. In particular, we did not vary them for any of our experiments, which included a diverse collection of real datasets. We therefore do not recommend users to change these thresholds.

## 3 Results

### 3.1 Experimental setup

#### Datasets

We used eight datasets, in order to test the robustness of isONclust with respect to different technologies, organisms, and read depths (Table 1). We first simulated three Iso-Seq read datasets from 107,844 unique cDNA sequences from ENSEMBL using SiMLOrD [41]. The datasets contained 100,000, 500,000, and 1,000,000 reads that were simulated with uniform distribution over the cDNA fragments. Next, we included a semi-biological Iso-Seq dataset (denoted RC0) where the transcripts are synthetically produced but then sequenced with Iso-Seq using the PacBio Sequel system. Then, we added three fully biological Iso-Seq datasets: PacBio Sequel datasets from a zebrafinch (ZEB) and a hummingbird (HUM), and a PacBio RSII system dataset from human brain tissue (ALZ). Finally, we included a ONT dataset of human cDNA sequenced with a MinION, which exhibits a different error profile and higher error rates than Iso-Seq. The non-simulated Iso-Seq and ONT datasets are publically available at [42] and [43], respectively.

**Table 1:** Datasets used for evaluation. The average error rate is computed from the quality score at each base of the reads (without homopolymer compression). A singleton class refers to a class that contains only one read. ^†^many of these originated from the synthetic spike-in non-human transcripts.

| Dataset | avg error rate (%) | n. classes |  | n. reads |  | % reads in non-singleton classes |
| --- | --- | --- | --- | --- | --- | --- |
|  |  | non-singleton | singleton | total | unaligned |  |
| ALZ | 1.7 | 13,350 | 10,187 | 814,667 | 98 | 98.7 |
| RC0 | 1.2 | 11,052 | 9,119 | 185,790 | 11,423<sup>†</sup> | 88.9 |
| HUM | 1.8 | 13,683 | 4,450 | 288,699 | 3,882 | 97.1 |
| ZEB | 1.9 | 12,891 | 4,936 | 309,749 | 129 | 98.4 |
| SIM-100k | 1.9 | 9,106 | 3,351 | 100,000 | 4 | 96.6 |
| SIM-500k | 1.9 | 14,792 | 2,152 | 500,000 | 4 | 99.6 |
| SIM-1000k | 1.9 | 16,510 | 1,594 | 1,000,000 | 4 | 99.8 |
| ONT | 12.9 | 14,863 | 13,665 | 890,503 | 38,061 | 94.2 |

#### Tools

The authors of carnac-lr [9] observed the inability of most clustering tools designed for other purposes [20, 25, 27, 14] to run on long-read transcriptomic data. However, we did consider four additional such tools: qCluster [30], linclust [18], DNACLUST [16], and MeShClust [15]. We also considered four tools specifically designed for long-read transcriptome data (carnac-lr, IsoCon, isoseq3-cluster, and Cogent). Isoseq3-cluster (which we will refer to simply as isoseq3) is the clustering tool used in the most recent version of PacBio’s *de novo* transcript reconstruction pipeline. Out of these eight tools, we found that only three (carnac-lr, isoseq3, and linclust) could process our two smallest datasets (SIM-100k and RC0). We therefore only include these tools in our final evaluations. All command lines and parameter settings are included in Appendix A.2.

#### Ground truth

Since the true clustering is not known, we use a clustering based on alignments to the reference genome as a proxy. We first align the reads with minimap2 [39] to the reference genome (hg38 for human, Tgut_diploid_1.0 for zebrafinch [44], and Canna diploid _1.0 for hummingbird [44]), with different parameters for Iso-Seq and ONT data (see Appendix A.2). The aligned reads are then clustered greedily by merging the clusters of any two reads whose alignments overlap. We refer to the cluster of a read obtained via this alignment to the reference as the *class* of the read. Reads that could not be aligned and hence could not be assigned to a class were excluded from all downstream accuracy evaluations. Some class metrics for the datasets are shown in Table 1.

Using alignments to the reference to define classes is an imperfect proxy of the true clustering. There are likely systemic misalignments due to gene sequence content, artifacts of the aligner, or chimeric reads due to *e.g.* reverse transcription errors. Thus our approach does not yield a reliable estimate for the absolute performance of a tool, but we believe it is a reasonable proxy to access the relative performance between different tools.

#### Evaluation metrics

There exists several metrics to measure quality of clustering. We mainly use the V-measure and its two components completeness and homogeneity [45]. Let *X* be an array of *n* integers, where *n* is the number of reads and the *i*^th^ value is the cluster id given by a clustering algorithm. Similarly, let *Y* be an array with the assigned ground truth class ids of the reads, ordered as in *X*. *Homogeneity* is defined as *h* = 1 *− H*(*Y |X*)*/H*(*Y*) and *completeness* as *c* = 1 *− H*(*X|Y*)*/H*(*X*). Here, *H*(*\**) and *H*(*\*|\**) refer to the entropy and conditional entropy functions, respectively [45]. Intuitively, homogeneity penalizes over-clustering, i.e. wrongly clustering together reads, while completeness penalizes under-clustering, i.e. mistakenly keeping reads in different clusters. The *V-measure* is then defined as the harmonic mean of homogeneity and completeness. These are analogous to precision, recall, and F-score measures for binary classification problems. We chose the V-measure metric as it is independent of the number of classes, the number of clusters, and the size of the dataset—and can therefore be compared across different tools [45]. Moreover, it can be decomposed in terms of homogeneity and completeness for a better understanding of the algorithm behavior.

In order to avoid bias with respect to a single accuracy measure, we also included the commonly used adjusted Rand index (ARI) [46]. Intuitively, ARI mesures the percentage of read pairs correctly clustered, normalized so that a perfect clustering achieves an ARI of 1 and a random cluster assignment achieves an ARI of 0. The formal definition is more involved [46] and, since it is standard, we omit it here for brevity.

In addition, we measure the percent of reads that are in non-singleton clusters. Since the coverage per gene is sufficiently high in all our datasets, the percentage of reads that are in non-singleton classes is high (89 - 100%, Table 1). Thus, any reads in singleton clusters in excess of this amount are indicative of reads that likely could have been clustered by the algorithm, but did not. Finally, we measure the runtime and memory usage (Table 2) of all the experiments.

**Table 2:**
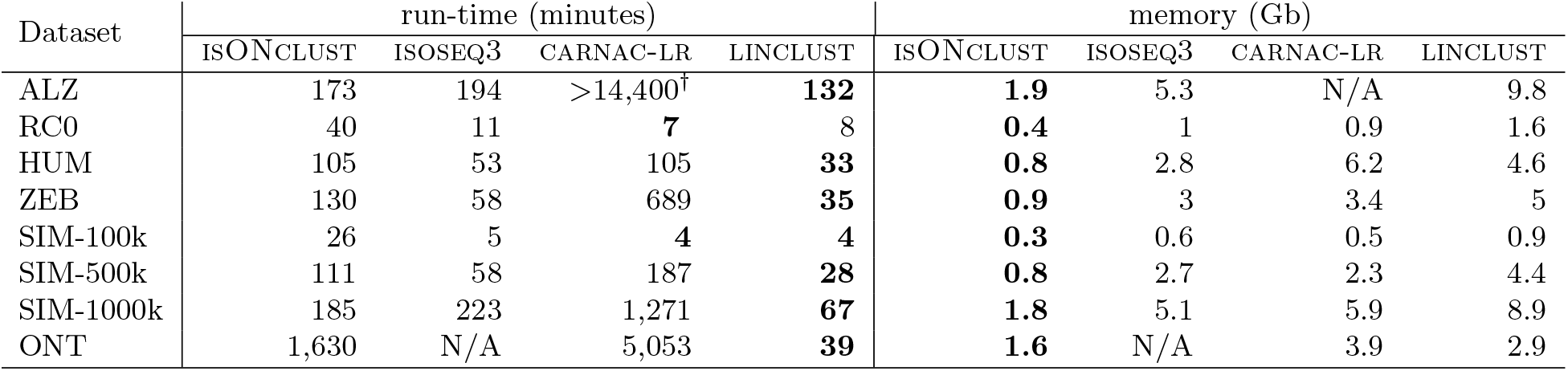
Run-time and peak memory usage for the clustering algorithms. isONclust was run on 1 core. The other tools were run with 8 cores specified. Runtime for carnac-lr includes mapping time with minimap. (^†^the run was terminated after 10 days.)

### 3.2 Comparison against other tools

The most direct comparison of our tool is to carnac-lr, which solves the same problem we do. One of its stated limitations is a worst-case cubic runtime [9], and we indeed observe that it does not scale well with growing sizes of our datasets (Table 2). For the largest Iso-Seq dataset (ALZ, 814k reads), carnac-lr did not complete within 10 days. For the other two large datasets (SIM-1000k and ONT), carnac-lr was *>* 6x and *>* 3x slower than isONclust, respectively. In terms of accuracy, carnac-lr performed reasonably well but always had a lower V-measure and ARI than isONclust. carnac-lr also placed less reads in non-singleton clusters than isONclust. For the ONT data, in particular, it was only able to place 54% of the reads into non-singleton clusters (compared to 94.5% for isONclust), even though 94.2% of the reads were in non-singleton classes (Table 1).

isoseq3 solves a slightly different problem than isONclust: its objective is to cluster reads together from the same isoform of a gene, rather than from the same gene family (*i.e.* in the case of alternative splicing, it will have separate clusters for each isoform). Thus, completeness, V-measure, and ARI with respect to our ground truth are not fair metrics by which to evaluate isoseq3. Nevertheless, isoseq3 leaves many reads unclustered: 26-36% of the reads from the real datasets and 53-89% of the reads from the simulated datasets (Table 3). In some cases, this could be caused by low coverage per isoform; however, SIM-1000k contains on average 9 reads per isoform, which should enable an algorithm to cluster substantially more than 53% of the reads. In terms of homogeneity, isoseq3 slightly outperforms isONclust, indicating that isoseq3 is the right tool if the goal is a conservative clustering. isoseq3 is designed for only Iso-Seq data and is thus not run on the ONT dataset.

**Table 3:** Performance and accuracy of the tools on our datasets. %NS is the percentage of reads in non-singleton clusters. The number of clusters is split between NS (non-singleton clusters) and S (singleton clusters).

| Dataset | Tool | accuracy |  |  |  | %NS<br>reads | n. clusters |  |
| --- | --- | --- | --- | --- | --- | --- | --- | --- |
|  |  | V | c | h | ARI |  | NS | S |
| ALZ | isONCLUST | **0.944** | **0.899** | 0.993 | **0.630** | **96.1** | 23265 | 32169 |
|  | ISOSEQ3 | 0.813 | 0.686 | **0.998** | 0.423 | 73.9 | 63512 | 212246 |
|  | LINCLUST | 0.839 | 0.725 | 0.996 | 0.518 | 80.8 | 57942 | 156476 |
| RC0 | isONCLUST | **0.977** | **0.961** | 0.994 | **0.804** | **90.1** | 12513 | 18459 |
|  | ISOSEQ3 | 0.923 | 0.859 | **0.997** | 0.640 | 66.6 | 14025 | 62085 |
|  | CARNAC-LR | 0.94 | 0.904 | 0.98 | 0.346 | 82.4 | 11002 | 32778 |
|  | LINCLUST | 0.933 | 0.877 | 0.996 | 0.566 | 77.7 | 18116 | 41363 |
| HUM | isONCLUST | **0.958** | **0.971** | 0.947 | **0.716** | **97.3** | 12140 | 7773 |
|  | ISOSEQ3 | 0.88 | 0.805 | **0.97** | 0.486 | 67.2 | 24171 | 94558 |
|  | CARNAC-LR | 0.934 | 0.944 | 0.924 | 0.489 | 93.3 | 9565 | 19323 |
|  | LINCLUST | 0.888 | 0.825 | 0.962 | 0.462 | 78.9 | 28066 | 61046 |
| ZEB | isONCLUST | **0.965** | **0.965** | 0.965 | **0.809** | **97.1** | 12767 | 8949 |
|  | ISOSEQ3 | 0.878 | 0.79 | **0.986** | 0.476 | 64.5 | 24097 | 110028 |
|  | CARNAC-LR | 0.93 | 0.94 | 0.92 | 0.401 | 93.4 | 9315 | 20555 |
|  | LINCLUST | 0.881 | 0.801 | 0.979 | 0.455 | 76.1 | 31119 | 74119 |
| SIM-100k | isONCLUST | **0.984** | **0.987** | 0.981 | **0.829** | **96.7** | 8931 | 3346 |
|  | ISOSEQ3 | 0.863 | 0.76 | **0.998** | 0.007 | 10.9 | 5013 | 89114 |
|  | CARNAC-LR | 0.979 | 0.99 | 0.969 | 0.734 | 96.1 | 8165 | 3945 |
|  | LINCLUST | 0.911 | 0.845 | 0.988 | 0.258 | 76.5 | 17856 | 23478 |
| SIM-500k | isONCLUST | **0.984** | **0.988** | 0.98 | **0.831** | **99.5** | 13996 | 2274 |
|  | ISOSEQ3 | 0.809 | 0.681 | **0.995** | 0.006 | 33.1 | 68704 | 334547 |
|  | CARNAC-LR | 0.971 | 0.974 | 0.967 | 0.695 | 97.1 | 12761 | 14527 |
|  | LINCLUST | 0.895 | 0.82 | 0.985 | 0.263 | 89.8 | 48608 | 51026 |
| SIM-1000k | isONCLUST | **0.984** | **0.988** | 0.98 | **0.832** | **99.8** | 15590 | 1945 |
|  | ISOSEQ3 | 0.788 | 0.654 | **0.993** | 0.006 | 46.8 | 180629 | 532410 |
|  | CARNAC-LR | 0.958 | 0.949 | 0.967 | 0.674 | 94.3 | 14423 | 56502 |
|  | LINCLUST | 0.89 | 0.813 | 0.984 | 0.264 | 91.8 | 68752 | 81641 |
| ONT | isONCLUST | **0.886** | **0.825** | 0.957 | **0.353** | **94.5** | 39464 | 48935 |
|  | CARNAC-LR | 0.797 | 0.669 | **0.984** | 0.095 | 54.2 | 27483 | 408270 |
|  | LINCLUST | 0.72 | 0.563 | 1 | <0.001 | 0.1 | 516 | 889346 |

Finally, we compare against linclust, which has a generic objective to cluster any sequences above a given sequence similarity and coverage. We explored several combinations of parameters to achieve the best results (see Appendix A.2). While linclust was the fastest tool, it has substantially worse accuracy on Iso-Seq data than other tools and was able to cluster only 0.1% of the ONT reads. This is not surprising, given that it was not designed for transcriptome data.

### 3.3 Performance observations

#### Scalability

For Iso-Seq, we can use the simulated data, which only varies in read depth, to conclude that isONclust has linear scaling with respect to the number of reads (Table 2). The absolute run-time is 3.1 hours for the largest Iso-Seq dataset, which is acceptable but could be further improved through parallelization or code optimization. For ONT data, the dearth of mature transcriptomic read simulators makes a controlled evaluation of scalability challenging. Though we are 3x faster than carnac-lr on our dataset, the absolute run-time is still fairly high (27.2 hours) and improving it is an immediate future goal. We expect that parallelization will yield significant speed-ups, keeping in mind that other tools were run on eight cores compared to only one core for isONclust (Table 2).

#### Role of class size

We investigated if isONclust’s clustering accuracy is affected by the class size. We binned the reads according to ranges of class size and computed the completeness and homogeneity with respect to each bin (Figure 1). The completeness clearly decreases with increased class size, indicating that isONclust tends to have more fragmented clusters as the class size increases. Homogeneity has no clear trend for class sizes up to 50, but decreases after that.

**Fig. 1:**
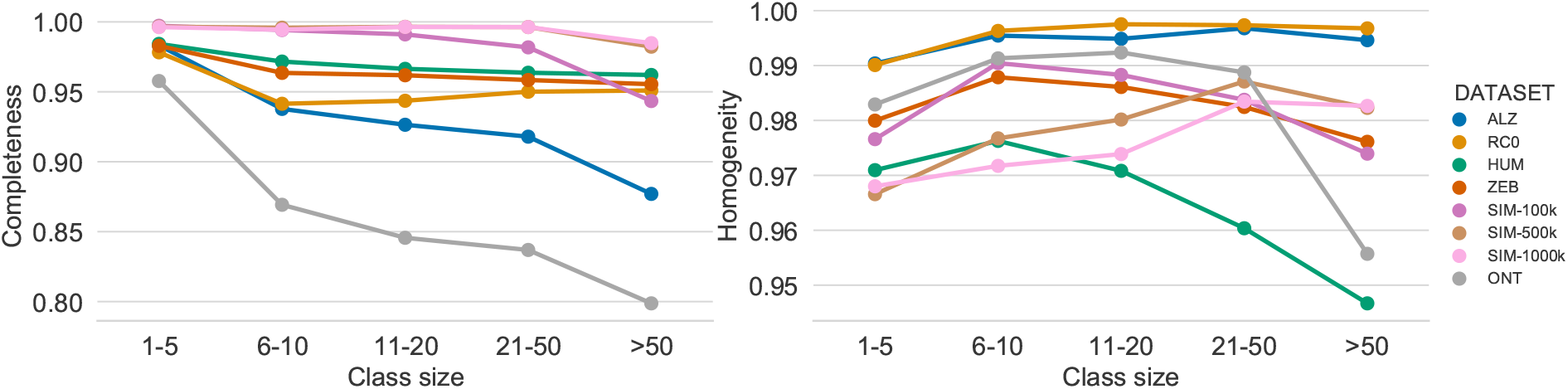
Completeness and homogeneity of isONclust across various class sizes.

#### Role of read error rates

Base errors pose a challenge to any clustering algorithm, so we measured how they affected our accuracy. We batch reads with respect to their error rate and measure the ARI within each batch (Figure 2, left panel). For Iso-Seq, isONclust has relatively stable ARI across different error rates (with ALZ being the exception), which we believe is due to our algorithm’s use of quality values. This is not true for ONT, where error rates of 7 - 20% have a detrimental effect on isONclust. Nevertheless, compared to carnac-lr, isONclust has substantially higher ARI across error rates, datasets, and technologies (Figure 2, right panel); e.g. for the ONT dataset, isONclust does better at 20% error rate than carnac-lr does at 7%.

**Fig. 2:**
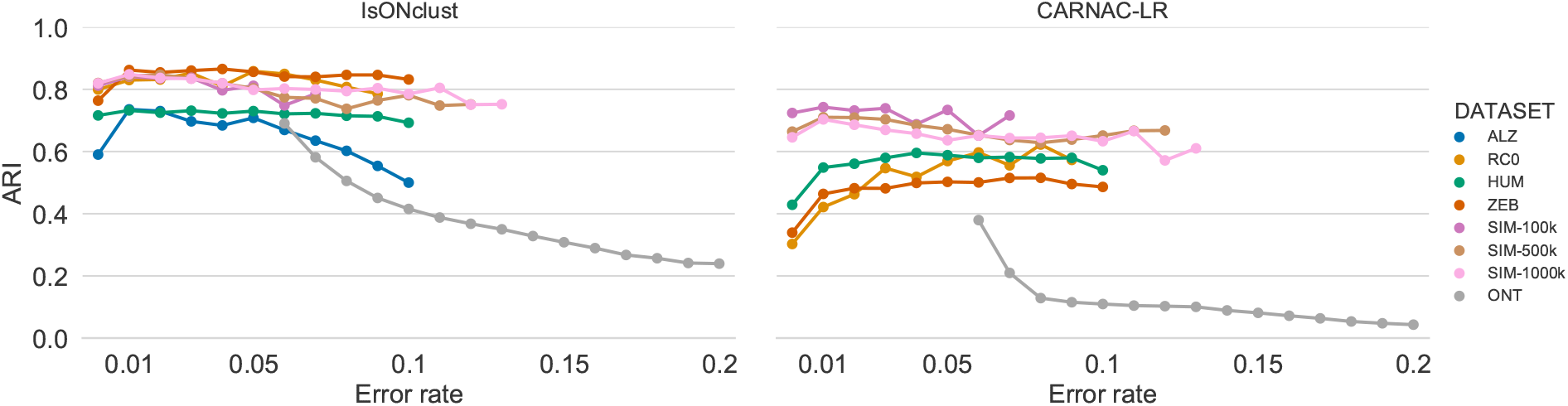
Accuracy (measured by the ARI) of isONclust and carnac-lr as a function of error rates. The read error rate is inferred by isONclust. Reads are binned according to their error rate, rounded to the nearest two decimal points. Datapoints for where there are at least 1,000 reads are shown.

#### Breakdown of algorithm stages

For each read, isONclust either assigns it to a new cluster or to an existing cluster. If the read goes to an existing cluster, then it is either by minimizer matching or by alignment. We measure the distribution of reads into these three cases for all our datasets (Figure 3). For the non-simulated Iso-Seq data, alignment was invoked only 6-10% of the time. However, for the ONT and simulated Iso-Seq data, alignment was invoked more frequently (22-34%), indicating room for future run-time improvement.

**Fig. 3:**
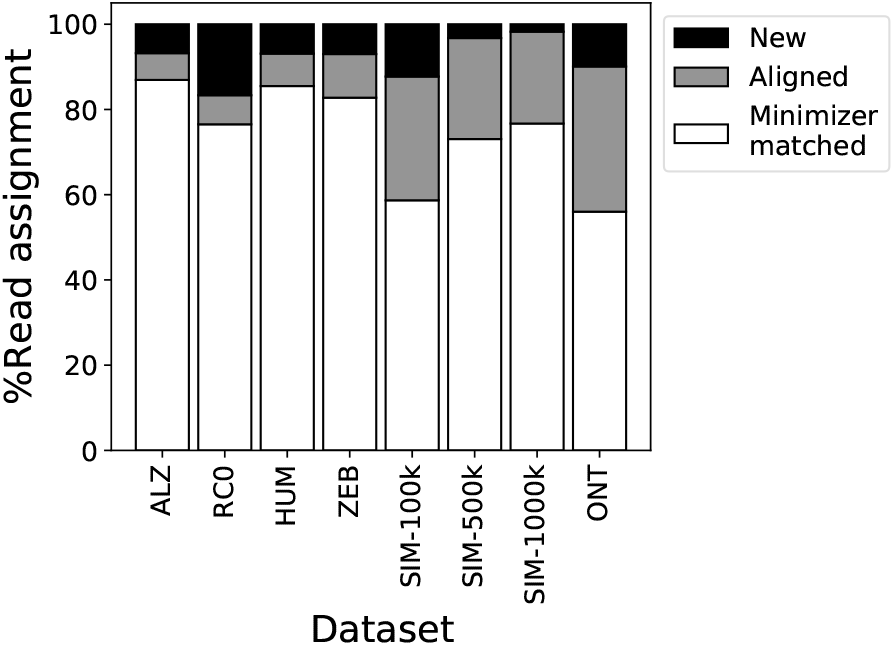
Distribution of the stages of our algorithm. A read is either minimizer-matcher or aligned to an existing cluster, or a new cluster is formed.

## 4 Conclusion

In this paper, we presented isONclust, a clustering tool for long-read transcriptome data. The design choices of our algorithm are mostly driven by scaling and the desire to use quality values. In order to scale, we made the algorithm greedy so that it can avoid doing an all-to-all similarity comparison. We avoid the natural but time consuming step of recomputing the best representative within a cluster after each update. Our initial sorting step mitigates the potentially negative effects of this by making sure that the representative is guaranteed to have the largest expected number of error-free *k*-mers among all reads in the cluster. Furthermore, we avoid the expensive alignment step whenever possible by using minimizer matching. In terms of quality values, we use them throughout the algorithm, including in the initial sorting step, in deciding whether mismatched minimizers are the result of sequencing error, and in computing pairwise alignment. The use of quality values is critical to the success of our algorithm and to its ability to handle both PacBio and Nanopore data.

Our results indicate that isONclust is a substantial improvement over existing methods, with higher accuracy and/or better scaling than other comparable tools. We also demonstrated that isONclust performs well across a breadth of instruments (PacBio’s Sequel, PacBio’s RCII, and Oxford Nanopore), organisms (human, zebrafinch, and hummingbird), with each also having a different quality of reference for estimating the ground truth, and read depths (from 100k to 1mil reads). In all these scenarios, isONclust outperforms others on all relevant accuracy metrics, with the exception that isoseq3 produces a more homogeneous clustering (though at the cost of clustering much fewer reads).

Ultimately, we would like to combine isONclust with a post-clustering error-correcting module in order to reconstruct transcripts *de novo* from non-targeted Iso-Seq and ONT data. We have previously taken this approach in our IsoCon tool [2] for targeted Iso-Seq data. IsoCon, however, is not able to scale to the much larger non-targeted datasets and to the higher error rates of ONT. With the development of isONclust, we are now able to overcome these challenges in the clustering step. Our next step is to tackle the error correction problem within each cluster. The ultimate goal is to develop a tool for *de novo* transcript reconstruction, which will be the first such tool for ONT data and an improvement over other methods for Iso-Seq data.

## Acknowledgements

This work has been supported in part by NSF awards DBI-1356529, CCF-551439057, IIS-1453527, and IIS-1421908 to PM.

### A Appendix

#### A.1 Alignment parameters

For Smith-Waterman alignment of a read *r* to a representative *c*, isONclust uses the following parameters: *match* = 2*, mismatch* = *−*2*, gapExt* = *−*1. *GapOpen* is set as a function of the combined error rate of *r* and *c*, denoted by *ϵ* = *ϵ_r_* + *ϵ_c_*: *gapOpen* = 2 for *ϵ >* 0.1, *gapOpen* = 3 for 0.04 *< ϵ ≤* 0.1, *gapOpen* = 4 for 0.01 *< ϵ ≤* 0.04, *gapOpen* = 5 for *ϵ ≤* 0.01.

#### A.2 Commands used for running tools

##### Carnac-LR

For carnac-lr we used version a5d8271d1bc503bcac00b615ee0673537ff99468 (git commit ID) and command line:

~~~
 $ minimap −Sw2 −L100 −t8 { input. flnc_fasta } { input . flnc_ fasta }
 $ python paf_to_CARNAC. py { minimap_out } { input . flnc fasta } { carnac_input }
 $ CARNAC−LR − f { carnac_input } −t 8 −o { carnac_output }
~~~

We used the same parameter for minimap as they did in their experiments [9].

##### isoseq3-cluster

For isoseq3, we used version sierra 0.7.1 (commit v0.4.0-121-g22a3096*) and command line:

~~~
 $ isoseq3 cluster ––num−threads 8 { input. ccs } { output. consensus }
~~~

##### isONclust

For isONclust with Iso-Seq data, we used:

~~~
$ isonclust. py ––t 8 ––flnc { input . flnc }
          –– ccs { input. ccs } ––outfolder { outfolder }
~~~

For the ONT data we ran isONclust as

~~~
 $ isonclust. py ––k 13 ––w 20 ––t 8 ––fastq { input . fastq }
    ––outfolder { outfolder }
~~~

##### Linclust

For linclust, we used version 822c8b57bb3ded9f37540b7cc2c9b97cf319d6e8 (the git commit ID) and command line:

~~~
 $ mmseqs easy− linclust ––seq−id−mode 1 ––cov−mode 1 ––threads 8
          ––kmer−per−seq [21, 100, 1000, 100000]
          −c [0.0, 0.4, 0.5, 0.6, 0.8 −e [0.1, 0.001]
          { input. flnc_fasta } { linclust output } { tmp_dir }
~~~

We ran linclust with default parameters, as well as with “–seq-id-mode 1 –cov-mode 1” and various combinations of “–kmer-per-seq”, “-c”, and “-e” after personal communication with author about suitable parameters for this type of data. In general we observed conservative results across all combinations. We present the results with the parameter setting performing the most permissive clustering “–seq-id-mode 1 –cov-mode 1 –kmer-per-seq 10000 -c 0.0 -e 0.1” as they in general gave the the best V-measure, completeness and percentage of non-trivially clustered reads without notably sacrificing homogeneity. The runtime and memory usage for these parameter settings were significantly higher than for other parameter settings.

##### qCluster

With qCluster we used the version available at website http://www.dei.unipd.it/ciomp-in/main/qcluster.html as of 10/23/1018 (no version number available) and ran

~~~
 $ q Cluster −d e −c [20000, 1000, 100] −k [15, 31] { input. reads }
     >{ output file }
~~~

We tried the parameter combinations within brackets but qCluster returned segmentation faults and seemed to occupy more than 264Gb of memory for all the combinations on our two smallest datasets SIM-100k and RC0.

##### MeShClust

With MeShClust we ran version 1.0.0 as follows

~~~
$ meshclust { input. reads } ––id [0.80, 0.90] ––threads 8
     ––output { output_file }
~~~

We tried the parameter combinations within brackets but we encountered a run-time error for all combinations that we tested on SIM-100k and RC0 (issue submitted https://github.com/TulsaBioinformaticsToolsmith/MeShClust/issues/6).

##### DNACLUST

With DNACLUST we ran release 3 as follows

~~~
 $ dnaclust { input. reads } −t 8 [−− left_gaps allowed]
     −s [0.8, 0.85, 0.9, 0.95] −k 5 ––approximate− filter > { output file }
~~~

We tried the parameter combinations within brackets but encountered segmentation fault 5 minutes in on the smallest simulated dataset SIM-100k and 3 hours in on RC0, but DNACLUST did not occupy more memory than what was available (264Gb).

##### Cogent

We ran Cogent version 3.3 according to tutorial on how to cluster large datasets (https://github.com/Magdoll/Cogent/wiki/Running-Cogent#running-family-finding-for-a-large-dataset). As the tutorial suggested, we ran precluster first, and then cogent on each cluster created by precluster separately. With cogent installed through conda, we ran:

~~~
   $ run preCluster. py cpus =8 #generates a folder “ precluster_out ”
   $ generate_batch_cmd_for_Cogent_family_finding. py ––cpus=8
     ––cmd filename=cmd_file_{ dataset name }
     pre Cluster. cluster_info. csv precluster_out
     { dataset_name }_cogent_out
 $ chmod +x cmd_file_ { dataset_name }
 $ . / cmd_file_ { dataset_name }
~~~

Cogents algorithm is however not suitable for large clusters generated by pre cluster, and will halt due to runtime complexity (personal communication with author). On the RC0 dataset we observed that one of the pre-clusters generated contained over 80,000 sequences. While we observed Cogent making progress on smaller clusters (< 100) it halted for the larger cluster (we killed the program after 72 hours).

##### IsoCon

We ran IsoCon v0.3.2.

`IsoCon pipeline – fl_reads < flnc . fastq > − outfolder </path/ to / output>`

IsoCon is not designed for nontargeted Iso-Seq or ONT data. The tool relies on exact start and end positions in transcript coming for the primer pairs designed for a targeted dataset.

##### minimap2

To align Iso-Seq reads to a reference genome we ran minimap2 with the following suggested parameters:

`minimap2 −t 8 −ax s p l i c e −uf −C5 { r e f } { output . f as tq } > { output . alignment`}

To align ONT reads to a reference genome we ran minimap2 with the following suggested parameters:

`minimap2 −t 8 −ax s p l i c e −uf −k14 { ref } { input . reads } > { output . alignment` }

